# AMMI and GGE biplot analysis of yield of different elite Wheat line under terminal heat stress and irrigated environment

**DOI:** 10.1101/2021.01.25.428058

**Authors:** Bishwas K. C., Mukti Ram Poudel, Dipendra Regmi

## Abstract

Wheat crop in Nepal faces terminal heat stress which accelerates the gain filling rate and shortens the filling period which leads to reduced grain weight, size, number, quality that is yield loss. For minimization of this loss, genotypic selection of high yielding lines should be performed understanding the gene-environment interaction. With the view to obtain a high yielding line with stable performance across the environments an experiment was conducted using 18 elite wheat line and 2 check varieties in alpha lattice design (2 replication and 5 blocks per replication) in different environments viz. irrigated and terminal heat stressed environment. The analysis of variance revealed that genotype, environment and their interaction had highly significant effect on the yield. Furthermore, which-won–where model indicated specific adaptation of elite lines NL-1179, NL-1420, BL-4407, NL-1368 to irrigated environment and BL-4919 and NL-1350 to terminal heat-stressed environment. Similarly, Mean-versus-stability study indicated that elite line BL-4407, NL-1368, BL-4919, NL-1350 and NL-1420 had above average yield and higher stability whereas elite lines Gautam, NL-1412, NL-1376, NL-1387, NL-1404 and N-1381 had below average yield and lower stability. Also, ranking elite lines biplot, PCA1 explaining 73.6% and PCA2 explaining 26.4% of the interaction effect, showed the rank of elite line, NL-1420 > NL-1368> NL-1350 > other lines, close to ideal line. From these findings, NL-1420 with high yield and stability can be recommended across both the environment while NL-1179 is adapted specifically for irrigated and NL-1350 adapted specifically for terminal heat-stressed environment.

## 1. INTRODUCTION

Wheat (*Triticum aestivum* L.) is believed to be cultivated first about 10000 years ago when hunting culture transited to agriculture. Nepal has 35 improved cultivars, 540 landraces and 10 wild relatives [1]. Wheat is important human food crop, ranks on top three cereal in the world because of its adaptability, nutritional value and high yield potential [2]. In Nepal, wheat occupies a major part of economy and ranks third major crop which is mainly used for bread and biscuits purpose [1]. Furthermore, it is an industrial crop because the grain along with stalk and chaff serves as industrial raw materials. Also, stalk and chaff are used as mulch, construction material and animal bedding [3].

Wheat has broad adaptation, however, it is most suited to temperate climate and high temperature can negatively impact the yield [4]. The risk of temperature is variable with stage of plant [3] such as optimum range of temperature for growth during sowing is 16 °C – 22 °C, for anthesis and grain filling is 12 °C – 22°C [5] while during the period of ripening is 21 °C – 25 °C [6]. Beyond these limits the production is effected, hereby the situation of global climate change and temperature rise is a major risk in wheat production system. This is because wheat plant exposed to temperature above 24 °C during anthesis and grain filling under goes terminal heat stress causing yield reduction, this reduction increases with longer exposure period [7].

In past 10 years, the cropping area of wheat has decreased from 731131 ha in 2009/10 to 703992 ha in 2018/19 that is around 4% decrease in cropping area. Furthermore, the increase in productivity has been in slow rates of 0.102 t/ha average increase per year in those 10 year. The productivity at start of decade (2009/10) was 2.13 ton/hectare reached to 2.85 ton/hectare by the end of decade (2018/19)[8]. In addition to this, the productivity of wheat in Nepal is low compared to that in world as shown clear in the following data: the worldwide productivity of wheat was 3.32 ton/ha in 2015, 3.42 ton/ha in 2016, 3.54 ton/ha in 2017 and 3.43 ton/ha in 2018 whereas Nepal had productivity of wheat 2.59 ton/ha in 2015, 2.33 in 2016, 2.55 ton/ha in 2017 and 2.76 ton/ha in 2018 [9].

As a consequence of the low productivity Nepal’s wheat import was 103705 tons in 2015, 217105 tons in 2016, 199626 tons in 2017 and 107467 tons in 2018 [9]. To minimize import it is necessary to increase the productivity of wheat because there is little scope for increasing wheat cultivated land in this region. Thus, focus should be given to break the yield barrier by genetic and development work and for increase in the production the constraint to wheat production: biotic and especially abiotic stresses should be mitigated [10].

Wheat contributes around 20% calories in the globe [9]. Around 85% of wheat is grown in developing countries where it seek to improve livelihood [11]. For this, improvement in yield is a most, currently improvement rates is 0.3 – 1.7% per annum for farm yield and 0.3 – 1.1% for the potential yield. The yield gap is between 26 % and 69% which gives a wide potential to raise yield through breeding and suitable intensification [12].

Global climate increase by 1.8 – 5.8 °C at the end of the century and has already changed the agronomic practice developed thousands of years ago [13].This gradual increment in temperature is shortening the wheat growing season [14]. Together with this the rainfall pattern has been altered such that there is need of strategies to moderate the effect of several biotic and abiotic stress to cope up with climate change effects [15]. Among these high temperature stress during reproductive development is termed as terminal heat stress [16].

A significant of part of the South Asian Region is under terminal heat stress including Nepal [1]. Heat stress causes multiple effect in wheat farming which include physiological effect (mainly chlorophyll deterioration and decreased leaf water), biochemical effect (especially reduced photochemical efficiency and stress metabolites accumulation), effect on growth and development (reduced growth duration and low leaf and tiller formation) leading to yield reduction (from quality to size to crop stand and seed development) [17]. These effect can be quantified as increase temperature by 1-2°C reduces grain mass which is mainly due to two reasons: accelerating grain growth rate and shortened grain filling period in wheat [18].

As a consequence, the heat exposure results in the decrease in grain weight [19] and loss of yield [20]. Finding has shown decrease of 1000 grain weight by around 67.3 % [21].Also, in late sown wheat yield can be from (25-35) % [22] to 47% [21]. This loss is rapidly increasing with time which can be exampled by two reports that is grain yield decline by 32 kg/ha [23] low loss which have climbed up to 1534 kg/ha (46%) [24] in late sown wheat. Thus, heat stress is a major predictor on the wheat yield globally [25] and in Nepal, with this it also has severe synergic effect to decrease wheat yield drastically when coincides with drought [26] towards end of growing season.

The simplest and common solution for heat stress is production of new cultivar that is genotype selection [27] [28] that is able to give stable yield in adverse environments too. To minimize the losses several improved varieties with high grain yield, stress tolerance and disease resistance have been developed but still problem exist in improving productivity and profitability of wheat farming. Thus, there is particular need to gain heat tolerance and develop heat tolerant new germplasm and technology through wheat breeding [22].

The genotype selection is dependent on the understanding of interaction among the genotypes, environment and crop management practices which can be characterized using statistical method [29]. The variety with highest average yield in all the test environment alone cannot be used for recommendation to the farmers; analysis of stability of variety in the environment that is G × E interaction and physiological basis are also to be studied [30]. Thus, stability analysis can be effective tool to select genotype for drought and heat tolerance.

This study is conducted to observe the yield stability of genotypes under heat stress condition. The genotype with low fluctuation of yield in stress and non-stress environment is suitable for this studies needs to find a stable genotype which gives constant yield focusing interaction of genotypes with environment and environmental stress such as heat; [31]. This study is conducted to observe the yield stability of genotypes under heat stress and non-stressed environments (irrigated).

## 2. METHODOLOGY

### 2.1 Field experimentation

The field experiment was conducted in two different environment viz. irrigated and terminal heat stress. The field in irrigated condition was sown in the last week of November to get the normal temperature to wheat crop in the reproductive and ripening stage. Delayed sowing, the last week of December, in the field of terminal heat stress condition was done to get higher temperature to wheat crop in the reproductive and ripening stage which causes heat stress.

### 2.2 Temperature and rain

The information of maximum and minimum temperatures recorded fortnightly; and total rainfall during each fortnight period was obtained from National Wheat Research Program, Bhairahawa and is presented in (Fig 1).

**Fig 1.**
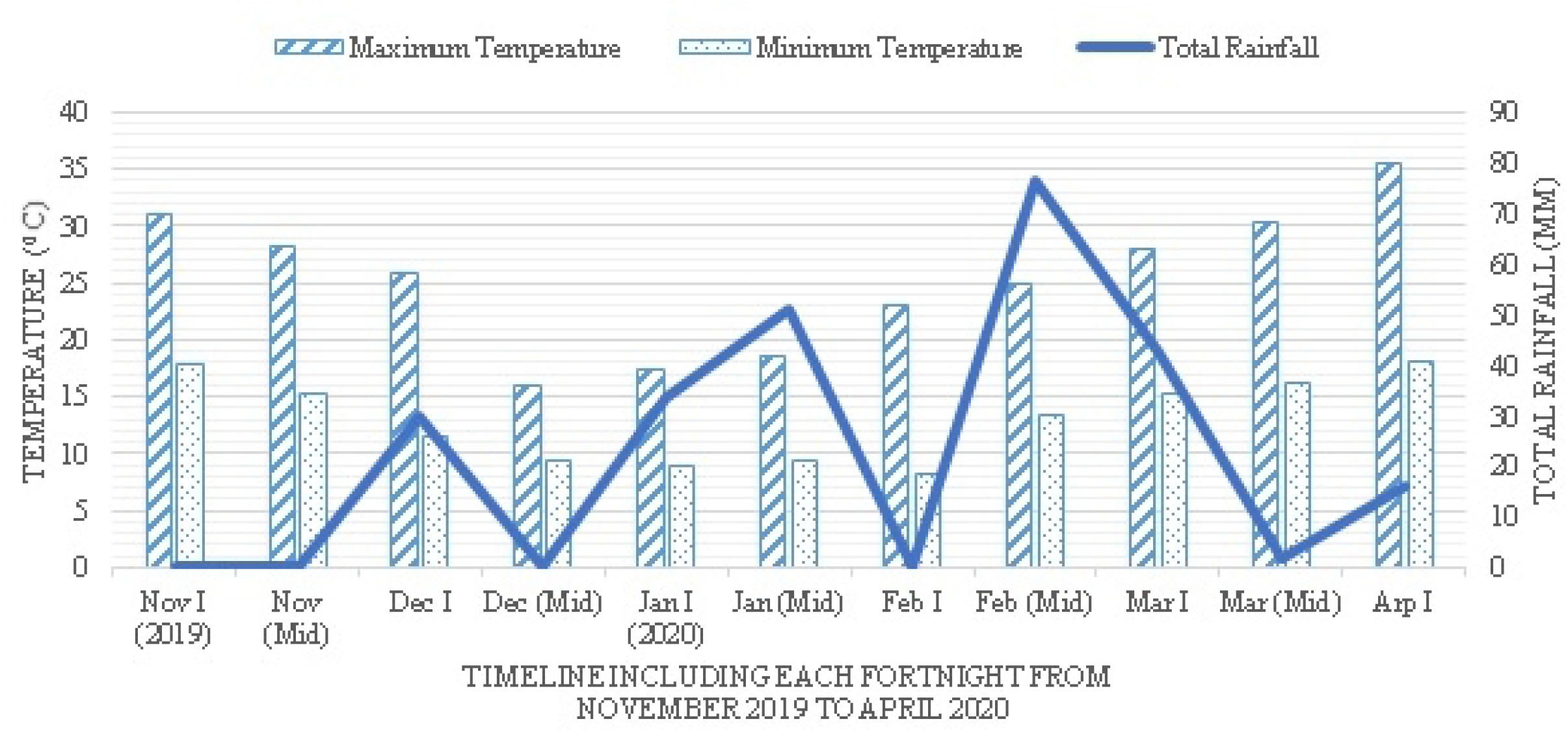
Maximum and minimum temperature; and total rainfall during November 2019 to April 2020 in the experiment filed

### 2.3 Soil properties

Soil samples obtained after land preparation was air-dried and well-grinded to sieve through 2mm sieve. Then soil characteristics analyzed in IAAS Soil Laboratory is given below:

Soil type: Clay loam
NPK content: 0.47 kg per ha (high) Nitrogen, 185 kg per ha (high) Phosphorus, 122.5 kg per ha Potassium
Organic matter content: 3.5%
Soil p^H^: 5.3 (acidic)

### 2.4 Plant materials

The research is conducted with 20 wheat genotypes collected from National Wheat Research Program, Bhairahawa which includes 15 Nepal Lines (NL), 3 Bhairahawa lines (BL) and two commercial varieties Gautam and Bhirkuti as check varieties. The complete set of genotypes with their entry name is given in (Table 1).

**Table 1.** List of elite wheat line with their origin, entry number as treatment.

| **Entry no.** | **Name of elite lines** | **Origin** | **Treatment** |
| --- | --- | --- | --- |
| 1. | Gautam | Nepal | T<sub>1</sub> |
| 2. | BL 4669 | Nepal | T<sub>2</sub> |
| 3. | NL1412 | CIMMYT, Mexico | T<sub>3</sub> |
| 4. | BL 4407 | Nepal | T<sub>4</sub> |
| 5. | NL 1368 | CIMMYT, Mexico | T<sub>5</sub> |
| 6. | NL 1417 | CIMMYT, Mexico | T<sub>6</sub> |
| 7. | Bhrikuti | CIMMYT, Mexico | T<sub>7</sub> |
| 8. | BL 4919 | Nepal | T<sub>8</sub> |
| 9. | NL 1376 | CIMMYT, Mexico | T<sub>9</sub> |
| 10. | NL 1387 | CIMMYT, Mexico | T<sub>10</sub> |
| 11. | NL 1179 | CIMMYT, Mexico | T<sub>11</sub> |
| 12. | NL 1369 | CIMMYT, Mexico | T<sub>12</sub> |
| 13. | NL 1350 | CIMMYT, Mexico | T<sub>13</sub> |
| 14. | NL 1420 | CIMMYT, Mexico | T<sub>14</sub> |
| 15. | NL 1384 | CIMMYT, Mexico | T<sub>15</sub> |
| 16. | NL 1346 | CIMMYT, Mexico | T<sub>16</sub> |
| 17. | NL 1404 | CIMMYT, Mexico | T<sub>17</sub> |
| 18. | NL 1413 | CIMMYT, Mexico | T<sub>18</sub> |
| 19. | NL 1386 | CIMMYT, Mexico | T<sub>19</sub> |
| 20. | NL 1381 | CIMMYT, Mexico | T<sub>20</sub> |

### 2.5 Experimental Design and Layout

The details of experiment was as follows:

Design: Alpha Lattice Design
Treatment Details: 20 treatments in 5 blocks each consisting of 4 treatments (in 4 plots)
Distance between any two blocks: 1 m
Distance between plots within a block: 0.5 m
Plot Area: 10 m^2^,

Dimension: 2.5 m × 4 m,
Sowing method: Continuous in a line
Number of rows: 10 rows
Row – row distance: 25 cm
Number of Replication (r) = 2
Number of Blocks (b) = 10
Number of blocks per replication (s) = 5
Number of treatments per block (k) = 4

### 2.6 Crop growth and management

The agronomic practices given below were followed:

Tillage: Ploughing followed by harrowing 1 week prior sowing; harrowing and leveling at sowing.

Fertilization: Farmyard Manure: 5 ton per ha

Recommended dose: NPK 100: 50: 25 kg per ha.
Terminal heat stress: Full dose at land preparation.
Irrigated: Half nitrogen and full dose P, K at land Preparation

Remaining half nitrogen at first irrigation.

Irrigation: 5 times each at CRI, Heading, Flowering, Milking and Soft dough Stage of wheat plant. Weeding: Manually at heading stage.

Harvesting: Manually using sickles when all maturity indices were complete.

Threshing: Manually using sticks.

Sample from 1m^2^ were kept separate form each plot for data collection of yield and related yield attributes.

### 2.7 Statistical analysis

MS Office 2013 was used data entry and processing. The AMMI Model with GGE bi-plots were used for analyzing the yield stability of elite lines in the heat stress and irrigated environment using GEAR (version4.0, CIMMYT, Mexico).

Additive Main Effect and Multiplicative Interaction (AMMI) model was used for the mean of yield of the 20 elite wheat lines from both the environments using GEA – R software. The AMMI model equation is:

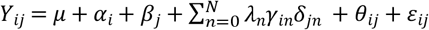

Where: Y_ij_ = the mean yield of elite line i in environment j, μ = the grand mean of the yield, α_i_= the deviation of the elite lines mean from the grand mean, β_j_ = the deviation of the environment mean from the grand mean, λ_n_ = the singular value for the PCA n, N = the number of PCA axis retained in the model, *γ_in_* = the PCA score of a elite line for PCA axis n, δ_jn_ = the environmental PCA score for PCA axis n, θ_ij_ = the AMMI residual and *ε_ij_* = the residuals. The degrees of freedom (DF) for the PCA axis were calculated based on the following method [32]. DF = G + E – 1 – 2n; Where: G = the number of elite lines, E = the number of environments and n = the n^th^ axis of PCA. The Genotype main effect plus Genotype by environment interaction (GGE) biplot used principal component comprised of set of elite lines scores multiplied by environment scores which gives a two dimensional biplot [33] and simultaneous study of the genotype plus genotype-environment interaction was performed.

## 3. RESULTS AND DISCUSSION

### 3.1 AMMI MODEL ANALYSIS

The result of analysis of variance of AMMI model revealed that grain yield is significantly (p < 0.001) affected by environment, genotype and genotype-environment interaction, which explained 75.66 %, 17.25% and 7.08 % of occurred variation, respectively. Furthermore, it showed two PCA with a highly significant (p < 0.001) different first interaction principal component of AMMI explaining 100% of the genotype environment interaction with 36 degree of freedom (df) that is 19 df of PCA1 and 17 df of PCA2 as shown in (Table 2).

**Table 2.** The analysis of variance of grain yield using AMMI models.

|  | **DF** | **SS** | **MS** | **F-value** | **PROB(F)** | **% explained** | **% accumulated** |
| --- | --- | --- | --- | --- | --- | --- | --- |
| **ENV** | 1 | 47244306 | 47244306 | 1516.25*** | 0 | 75.66 | 75.66 |
| **GEN** | 19 | 10773431 | 567022.7 | 18.2*** | 0 | 17.25 | 92.92 |
| **ENV\*GEN** | 19 | 4423810 | 232832.1 | 7.47*** | 0 | 7.08 | 100 |
| **PCA1** | 19 | 4316190 | 227167.9 | 7.61*** | 0 | 100 | 100 |
| **PCA2** | 17 | 0 | 0 | 0 | 1 | 0 | 100 |
| **Residuals** | 40 | 1246350 | 31158.75 | NA | NA | 0 | 0 |

ENV – Environment, GEN – Genotype (elite wheat lines) PCA – Principal Component of AMMI, DF – Degree of Freedom, SS – Sum of Square, MS – Mean Sum of Squares,

The AMMI biplot has the main effect as grain yield in the abscissa and the PCA1 as the ordinate where the genotypes or environment which lies on the same vertical line have same yield and which lies on same horizontal line have same interaction pattern. Also, the vectors of genotypes which have PCA1 close to origin (zero) has general adaptability whereas the vectors with larger PCA1 are specifically adapted to an environment.

In the AMMI biplot as shown on (Fig 2), the genotypes that cluster together behaves similar across the environment. The elite wheat lines: 5 (NL 1368), 8 (BL 4919), 13 (NL 1350), 14(NL 1420) are cluster close which performs similar in both terminal heat stress and irrigated environment. The heat stressed environment (2) have lower than average yield and irrigated environment have higher than average yield. The elite wheat line 18 is the most stable among the tested line and line 4 (BL 4407), 8 (BL 4919), 10 (NL 1387), 12 (NL 1369), 16 (NL 1346) are relatively stable lines in yield that is broadly adapted lines. The elite wheat lines 2 (BL 4669), 3 (NL 1412), 6 (NL 1417), 7 (Bhrikuti), 11 (NL 1179), 19 (NL 1386) are relatively unstable lines in yield because these lines are far from origin and can be specifically adapted to an environment. Specially, line 11 (NL 1179) are specifically adapted to irrigated environment and lines 1 (Gautam), 17 (NL 1404), 20 (NL 1381) are specifically adapted to terminal heat stressed environment.

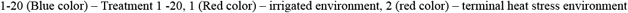

**Fig 2.**
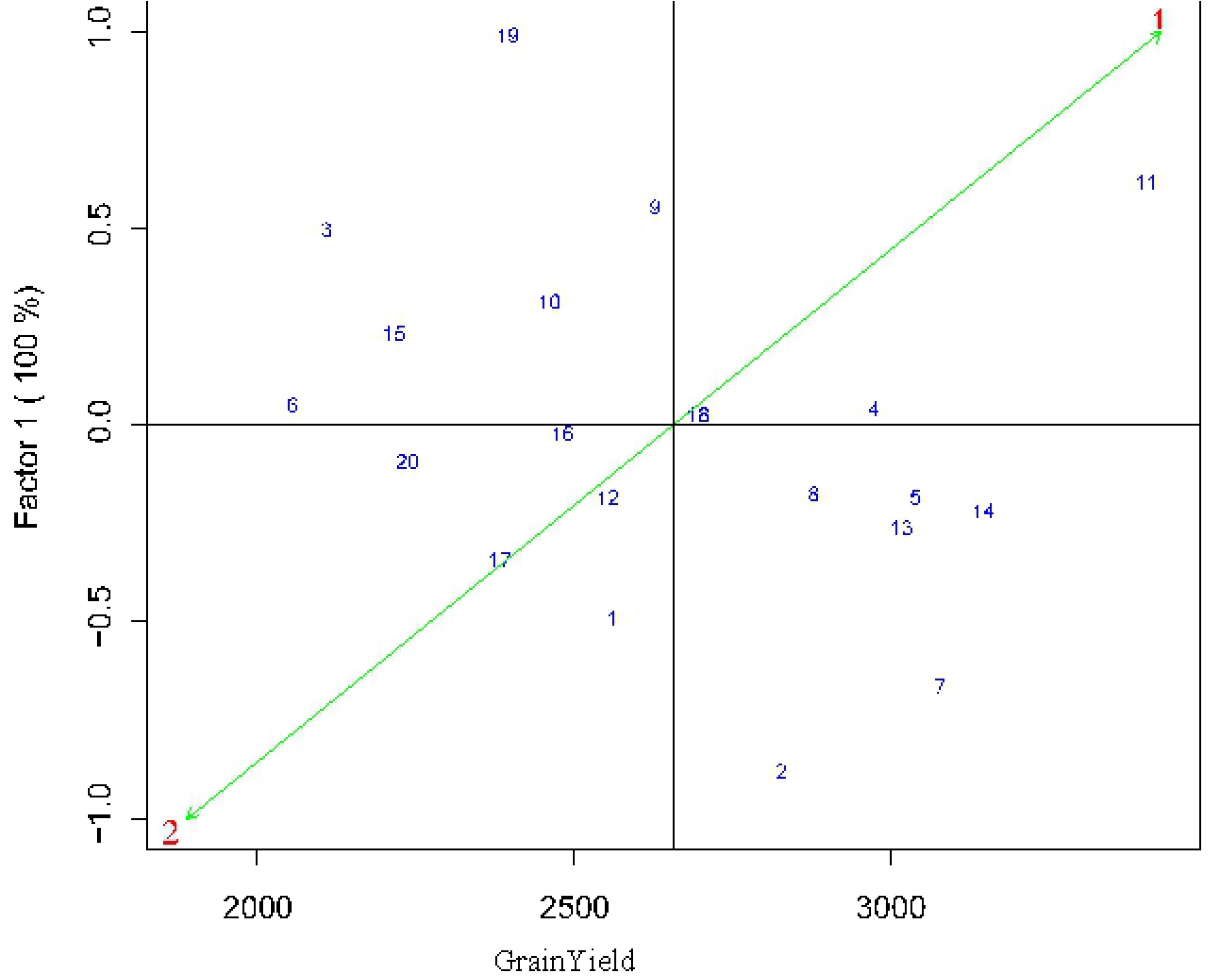
AMMI biplot PCA 1 versus grain yield of 20 elite wheat lines in terminal heat stress and irrigated environment

Similarly, the PCA 1 and PCA 2 scores is reported as representation of the stability of the lines across the environment that is the lines with the least PCA scores have high stability and vice-versa. According to the PCA1 score, line 2 (BL 4669)with score of −0.873 is the most stable followed by line 7 (Bhrikuti), 1 (Gautam) with score of −0.653, −0.481 respectively while PCA2 score shows lines 12 (NL 1369), 19 (NL 1386),9 (NL 1376) with score of −2.28 × 10^-9^, −1.41 × 10^-9^, −7.92 × 10^-10^ respectively are the most stable lines with regards to yield across both the test environments as shown in the (Table 3).

**Table 3.** Interaction Principal Component of AMMI (IPCA) 1 and 2 with yield of 20 test elite wheat lines.

|  | NAME | YLD | IPCA1 | IPCA2 |
| --- | --- | --- | --- | --- |
| 1 | 1(Gautam) | 2560.5 | -0.481 | $3.46 \times 10^{-09}$ |
| 2 | 10 (NL 1387) | 2463.25 | 0.324 | $-4.58 \times 10^{-10}$ |
| 3 | 11 (NL 1179) | 3405.25 | 0.627 | $9.24 \times 10^{-09}$ |
| 4 | 12 (NL 1369) | 2555 | -0.174 | $-2.28 \times 10^{-09}$ |
| 5 | 13 (NL 1350) | 3018.25 | -0.254 | $3.58 \times 10^{-10}$ |
| 6 | 14 (NL 1420) | 3146.75 | -0.208 | $2.93 \times 10^{-10}$ |
| 7 | 15 (NL 1384) | 2217.5 | 0.240 | $-3.39 \times 10^{-10}$ |
| 8 | 16 (NL 1346) | 2484 | -0.009 | $1.31 \times 10^{-11}$ |
| 9 | 17 (NL 1404) | 2384.25 | -0.330 | $4.66 \times 10^{-10}$ |
| 10 | 18 (NL 1413) | 2698.5 | 0.036 | $-5.11 \times 10^{-11}$ |
| 11 | 19 (NL 1386) | 2398.25 | 1 | $-1.41 \times 10^{-09}$ |
| 12 | 2 (BL 4669) | 2828.5 | -0.873 | $1.23 \times 10^{-09}$ |
| 13 | 20 (NL 1381) | 2239.75 | -0.082 | $1.16 \times 10^{-10}$ |
| 14 | 3 (NL 1412) | 2111.25 | 0.505 | $-7.13 \times 10^{-10}$ |
| 15 | 4 (BL 4407) | 2976.25 | 0.050 | $-7.16 \times 10^{-11}$ |
| 16 | 5 (NL 1368) | 3040.5 | -0.172 | $2.44 \times 10^{-10}$ |
| 17 | 6 (NL 1417) | 2057 | 0.058 | $-8.20 \times 10^{-11}$ |
| 18 | 7 (Bhrikuti) | 3079.75 | -0.653 | $9.22 \times 10^{-10}$ |
| 19 | 8 (BL 4919) | 2880.5 | -0.165 | $2.33 \times 10^{-10}$ |
| 20 | 9 (NL 1376) | 2629.5 | 0.561 | $-7.92 \times 10^{-10}$ |

PCA 1 score revealed that lines 19 (NL 1386), 11 (NL 1179), 9 (NL 1376), 3 (NL 1412) are relatively unstable line with scores of 1, 0.627, 0.561, 0.505 respectively while PCA 2 score shows that lines 11 (NL 1179), 1(Gautam), 2 (BL 4669) with score of 9.24 × 10^-09^, 3.46 × 10^-09^, 1.23 × 10^-09^ are relatively unstable lines with regards to yield across both the test environments.

The interaction principal component of AMMI (1 & 2) with yield of the 20 elite wheat lines are as follows:

### 3.2 GGE BIPLOT ANALYSIS

#### 3.2.1 Which Won Where Model

The most effective and succinct way of summarizing the genotype and genotype environment interaction of the dataset is the polygon-view of GGE biplot which visualizes the which-won-where pattern of an multi-environment dataset (Yan and Kang,). The polygon is drawn by joining the markers located farthest from the origin such that all other markers are included within the polygon.

The polygon view of this experiment as shown in the (Fig 3) revealed the 20 elite wheat lines fall under 6 sector and 2 test environment fall under 2 sectors in the polygon. The sector with irrigated environment consist of elite wheat lines: 4 (BL 4407), 5 (NL 1368), 11 (NL 1179) and 14 (NL 1420); indicating these lines are responsive in this environment. The elite wheat line 11 (NL 1179) vector is characterized by longest distance from the origin and is the vertex line of the sector implies line 11 (NL 1179) with specific adaptation in irrigated environment but lower stability in overall environment. Likewise, sector with terminal heat stressed environment consists of elite wheat lines: 8 (BL 4919) and 13 (NL 1350). The line 13 (NL 1350) vector had the relatively longer distance compared to line 8 from the origin indicating this as most responsive line in terminal heat stressed environment. Also, line 8 (BL 4919) and 13 (NL 1350) have higher stability because the distance of line vector was short from the origin.

**Fig 3.**
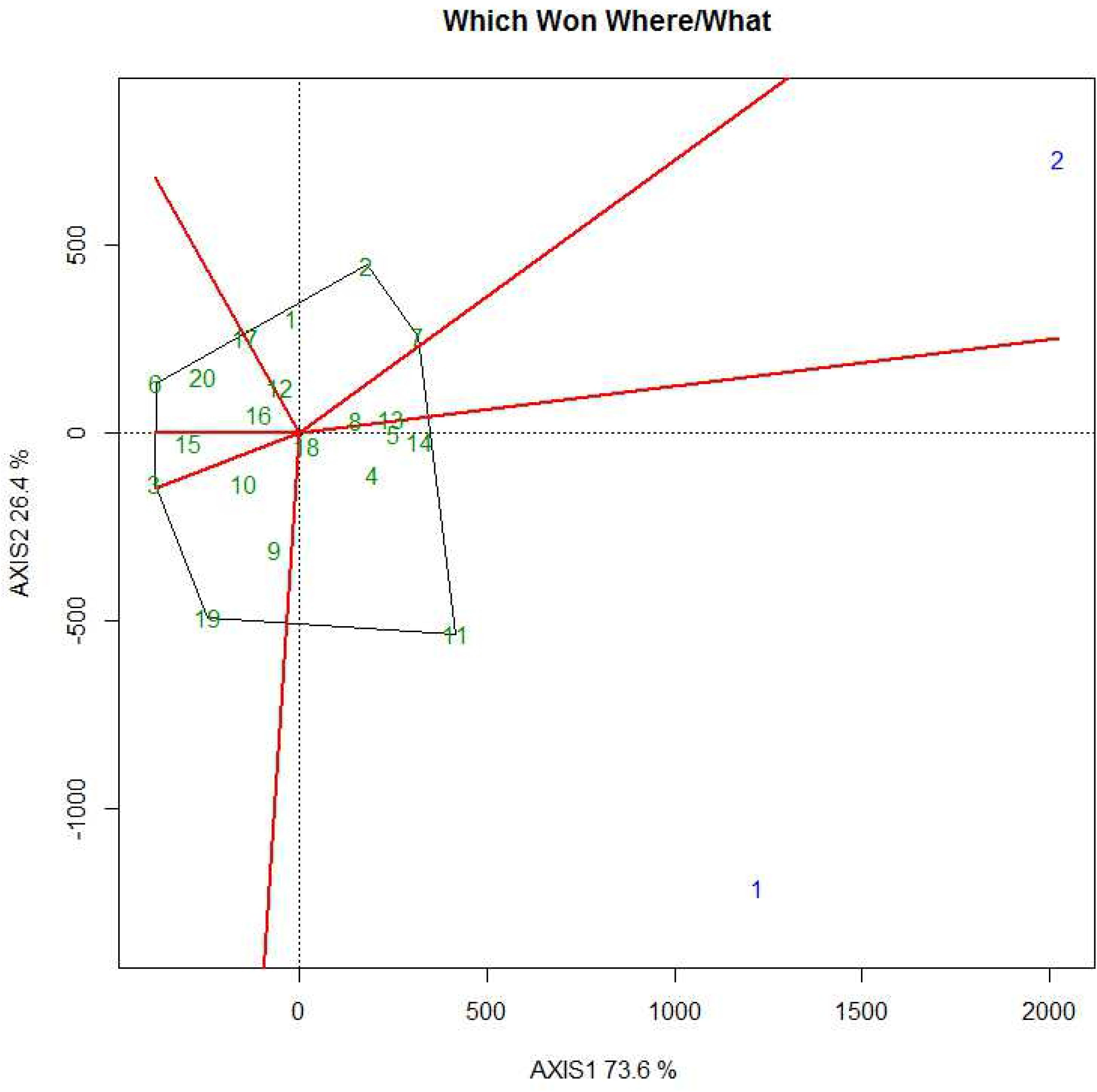
Polygon view of GGE biplot (which-won-where model) showing 20 elite wheat line in irrigated and terminal heat stressed environment

Thus, the which-won-where pattern of the trail revealed line 11 (NL 1179) as wining line in irrigated environment while line 13 (NL 1350) as wining line in the heat stressed environment. In addition, the polygon view showed elite wheat line 18 (NL 1413) near the origin of the biplot which means this line ranks the same in both test environments and lines: 8 (BL 4919), 12 (NL 1369) and 16 (NL 1346) are more stable. Also, elite wheat lines: 1(Gautam), 2 (BL 4669), 3 (NL 1412), 6 (NL 1417), 7 (Bhrikuti), 9 (NL 1376), 10 (NL 1387), 12 (NL 1369), 15 (NL 1384), 16 (NL 1346), 17 (NL 1404), 19 (NL 1386), 20 (NL 1381) are present in sector with no test environment symbolizes these lines are poor adapted to both the environments.

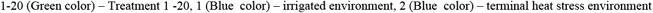

#### 3.2.2 Mean vs. Stability

When which –won-where pattern suggested wining elite wheat lines in the environments. There is need analyze mean performance and stability of all the elite wheat lines to make selection decisions. GGE biplot visualize performance and stability graphically with the help of Average Environment Coordinates (AEC). AEC is the mean of first and second principal components scores of the test environments which is represented by arrowhead in the (Fig 4). The line passing through arrowhead and origin is AEC abscissa and line perpendicular to it at origin is ordinate.

**Fig 4.**
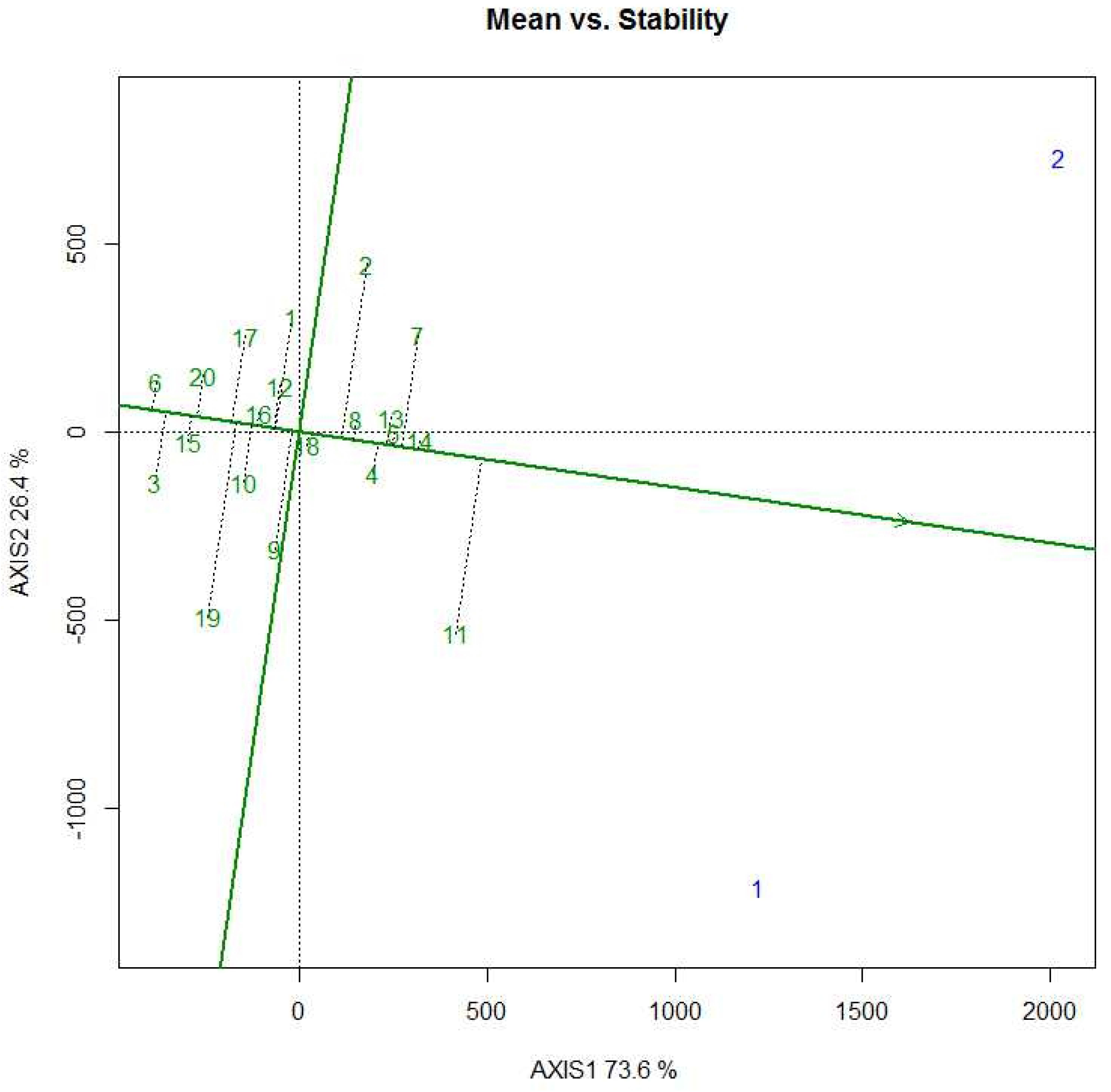
Mean vs. Stability view of GGE biplot showing the mean performance and stability of 20 elite wheat line in irrigated and terminal heat stressed environment.

Length of Abscissa gives the yield of genotypes that is above average and below average yield if right and left of the origin respectively, and length of ordinate approximate the GEI associated with the genotype that is more length corresponds higher variability and lower stability and vice-versa.

(Fig 4) shows elite wheat lines: 4 (BL 4407), 5 (NL 1368), 8 (BL 4919), 13 (NL 1350) and 14 (NL 1420) are above average yielders with more stability whereas lines: 2 (BL 4669), 7 (Bhrikuti), 10 (NL 1387) are above average yielders but with lower stability. Moreover, lines 6 (NL 1417), 12 (NL 1369) 15 (NL 1384), 16 (NL 1346), 20 (NL 1381) are stable but are below average yielders and lines 1(Gautam), 3 (NL 1412), 9 (NL 1376), 10 (NL 1387), 17 (NL 1404), 19 (NL1381) are both below average yielders with low stability.

Ideal lines have highest yield and absolute stability lying in the arrowhead and distance of other lines measures the desirability of lines. (Fig 4) show low desirability of these lines, however line 11 (NL 1179) unstable and 14 (NL 1420) stable are comparatively more desirable than other tested lines. These desirability gives the lines ranks in order line: 11 (NL 1179) followed by 14 (NL 1420), 7 (Bhrikuti), 5 (NL 1368), 13 (NL 1350), 4 (BL 4407) and as shown in (Fig 4).

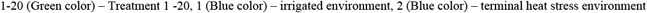

#### 3.2.3 Ranking Elite Wheat Lines (Genotypes)

The ideal line which practically not possible lies in the arrow head. To rank the lines coordinate is drawn:-line joining arrowhead and origin: first axis and line perpendicular to it at origin: second axis and the concentric circles with the arrowhead aids in further ranking as per the distance from arrowhead in the ordinate and inclusion in the circles.

The line 14 (NL 1420) is very close to the ideal line which can be used as reference in the lines evaluation. This is followed by lines 5 (NL 1368), 13 (NL 1350), 4 (BL 4407), 8 (BL 4919) in the rank of desirable genotypes which could be used for further testing in the heat stress and non-stress environments as shown in (Fig 5). The general ranking from the biplot is as follows:

**Fig 5.**
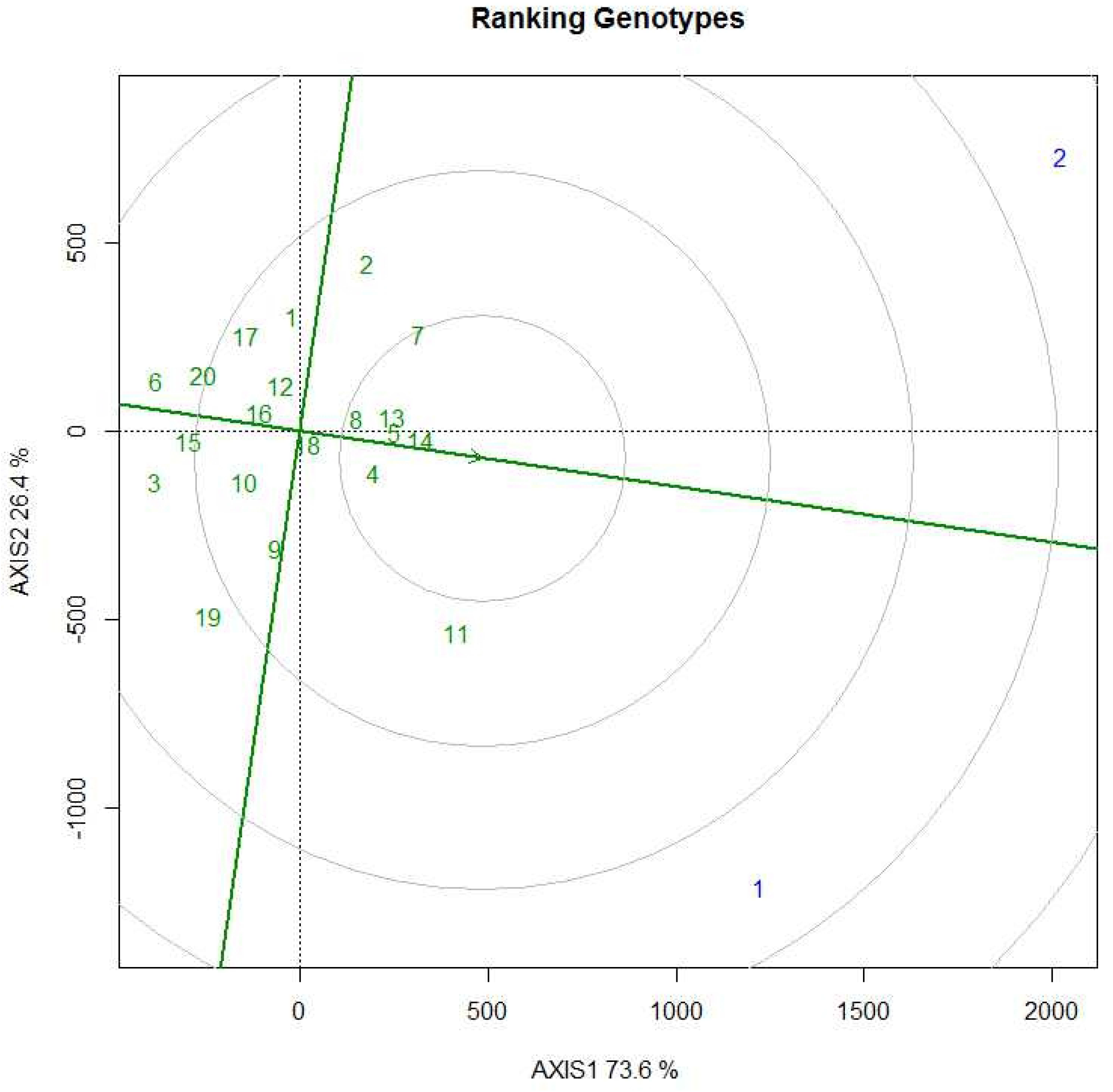
GGE biplot showing ranking of 20 elite wheat line with reference to ideal line in irrigated and terminal heat stressed environment

14 (NL 1420) > 5 (NL 1368) > 13 (NL 1350) > 4 (BL 4407) > 8 (BL 4919) >7 (Bhrikuti) > 11 (NL 1179)> 2 (BL 4669) > 18 (NL 1413) > 9 (NL 1376) > 12 (NL 1369) > 1(Gautam) > 16 (NL 1346) > 10 (NL 1387) > 17 (NL 1404) > 20 (NL 1381) > 15 (NL 1384) > 19 (NL 1386) > 3 (NL 1412) > 6 (NL 1417).

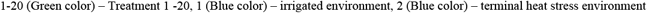

The comparison of biplot ranking and mean yield ranking of the genotypes in the combined environment (terminal heat stress and irrigated environment) is given in the (Table 4).

**Table 4.** Comparison of rank of 20 elite wheat lines based on mean yield and biplot ranking.

| Genotype Rank | Mean Yield Ranking | Biplot Ranking |
| --- | --- | --- |
| 1 | NL1179 | NL1420 |
| 2 | NL1420 | NL1368 |
| 3 | Bhirkuti | NL1350 |
| 4 | NL1368 | BL4407 |
| 5 | NL1350 | BL4919 |
| 6 | BL4407 | Bhirkuti |
| 7 | BL4919 | NL1179 |
| 8 | BL4669 | BL4669 |
| 9 | NL1413 | NL1413 |
| 10 | NL1376 | NL1376 |
| 11 | Gautam | NL1369 |
| 12 | NL1369 | Gautam |
| 13 | NL1346 | NL1346 |
| 14 | NL1387 | NL1387 |
| 15 | NL1386 | NL1404 |
| 16 | NL1404 | NL1381 |
| 17 | NL1381 | NL1384 |
| 18 | NL1384 | NL1386 |
| 19 | NL1412 | NL1412 |
| 20 | NL1417 | NL1417 |

#### 3.2.4 Discriminativeness vs. representativenss

The environment with no discriminating ability gives no information of lines that is useless and environment not representative is useless as well as misleading.

The GGE biplot use the vector of the environment to measure discriminitiveness that is more the length of environment vector more is the standard deviation within the environment indicating higher discriminating ability. The heat stress environment vector has comparatively more length ensuring it has a higher discriminating ability as shown in (Fig 6). Furthermore, cosine of angle between the environment gives the interrelationship between the environment that is angle just less than 90 ° shows a positive but low correlation coefficient between terminal heat stress and irrigated environment. Since the angle is large the environment are not redundant.

**Fig 6.**
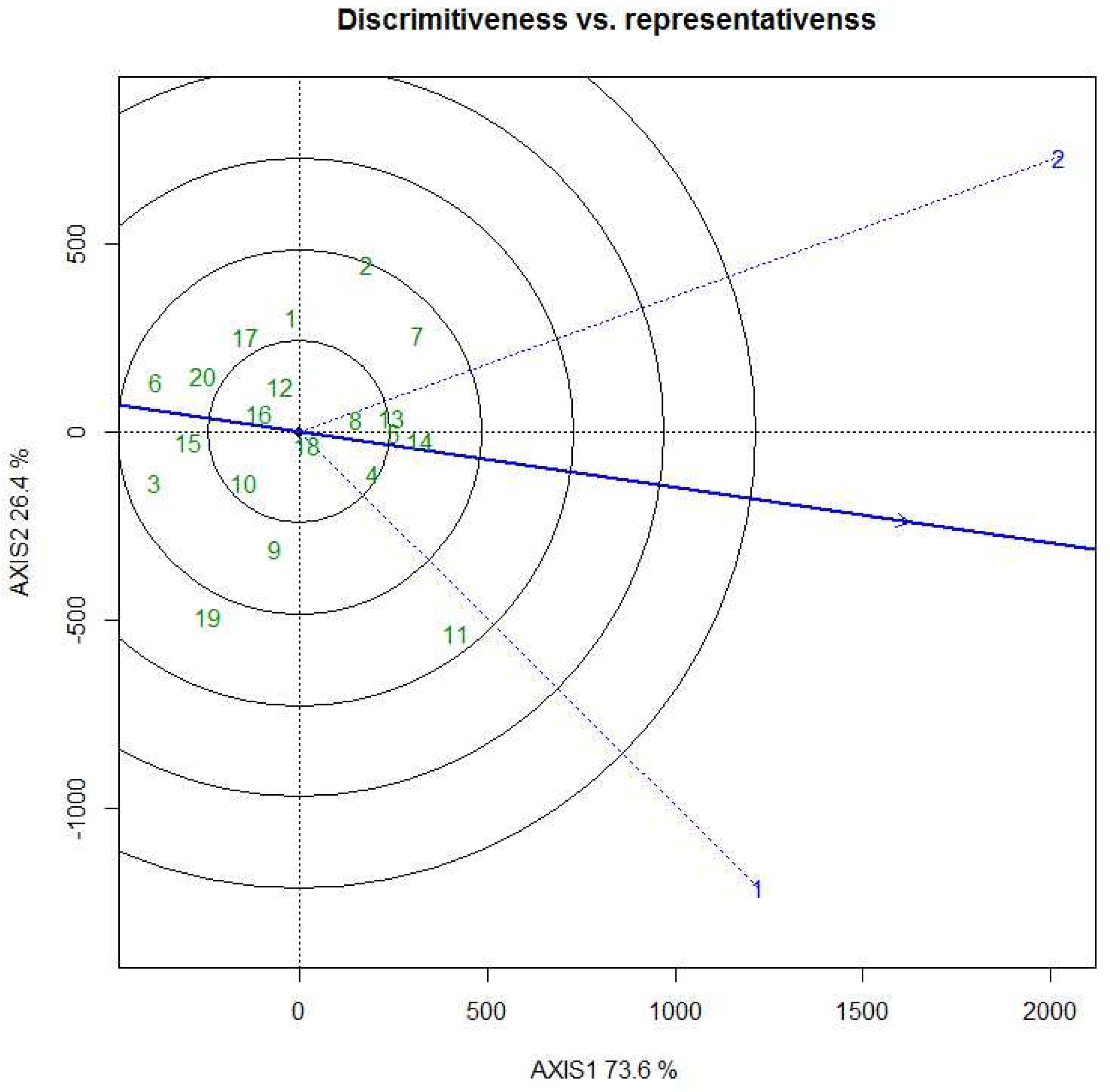
Discriminativeness vs. representativeness view of GGE biplot showing 20 elite wheat lines in irrigated and terminal heat stressed environment

Representativeness is measure of environment similar to the AEC ranking of genotypes. The desirability of environment is not clearly seen because of use of few environments. But, both the environment vector inscribe somewhat equal angle to the average environment coordinate symbolizes similar representativeness of irrigated and terminal heat stressed environment as shown in (Fig 6).

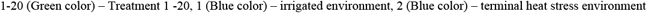

## 4. CONCLUSION

This study indicated that genotype, environment and their interaction have significant effect on the yield and the 100% of the interaction effect was explained by PCA 1 as per the AMMI model. Further analysis through GGE biplot concluded elite wheat line NL 1179 was specifically adapted to the irrigated environment whereas elite wheat line NL 1350 was specifically adapted to the terminal heat stressed environment via. Which-won where model. However, NL 1179 was among the least stable lines along with BL 4669 and NL 1386. The study of the mean vs stability and ranking of the line in the GGE biplot revealed elite wheat line NL 1420 along with NL 1368, NL 1350, BL 4919 were among stable line across the environment with higher than average yield. The terminal heat stressed environment had slightly higher discriminating ability than irrigated environment with comparatively equal representativeness. All in all, NL 1420 can be used for breeding programs as stable, high yielding line and for farmers NL 1179 and NL 1350 can be used for high yield with adaptability in irrigated and heat stressed environment respectively.

## 7. Acknowledgment

We are thankful to Pakhlihawa Campus, Institute of Agriculture and Animal Science, Rupandehi; Nepal Wheat Research Program, Rupandehi and Agri-botany Division, Nepal Agricultural Research Council, Kathmandu for their material support, valuable suggestion and facilitation for the field work which contributed a lot in successful completion of the research work.

## References

1. Joshi BK, Mudwari A, Bhatta M. Wheat Genetic Resource in Nepal. Nepal Agriculture Resource Journal. 2006; 7. doi: 10.3126/narj.v7i0.1859

2. Shewry PR. Wheat: Darwin review. Journal of Experimental Botany. 2009; 60(6): 1537–1553. doi:10.1093/jxb/erp058

3. Oyewole CI. The wheat crop.2016. Available from: https://www.researchgate.net/publication/310458715

4. Olabanji OG, Omeje MU, Mohammed I, Ndahi WB, Nkema I. Wheat. In: Cereal Crops of Nigeria: Principles of Production and Utilization. 2007; 22(337): 230–249.

5. Farooq M, Bramley H, Palta JA, Siddique KHM. Heat stress in wheat during reproductive and grain-filling phases Crit Rev Plant Sci. 2011; 30: 491–507. doi: 10.1080/07352689.2011.615687

6. Flato G, Marotzke J, Abiodun B, Braconnot P, Chou SC, Collons W, Cox P, Driouch F, Emori S, Eyring V, Forest C, Gleckler P, Guilyardi E, Jakeob C, Kattsov V, Reason C, Rummukainen M. Evaluation of Climate Models Climate change 2013, The Physical Basis, Working Group I Contribution to the fifth Assessment Report of the Intergovernmental Panel on Climate change. Cambridge University Press. 2013: 741–882. doi: 10.1017/CBO9781107415324.020.

7. Prasad PVV, Djanaguiraman M. Response of floret fertility and individual grain weight of wheat to high temperature stress: sensitive stages and thresholds for temperature and duration. Functional Plant Biology. 2014; 41:1261–1269.

8. MoALD. Area, Production and Yield by Major Cereal Crops, Last Ten Years. 2020. Available from: https://s3-ap-southeast-1.amazonaws.com/prod-gov-agriculture/server-assets/publication-1568704754479-585b7.xlsx

9. FAOSTAT. Online statistical database: Food and agriculture data. FAOSTAT. 2020. Available from: https://www.fao.org/faostat/en/#data/FS

10. Chatrath R, Mishra B, Ortiz-Ferrara G, Singh SK, Joshi A. Challenges to wheat production in South Asia. Euphytica. 2007; 157: 447–456. doi:10.1007/s10681-007-9515-2.

11. CAS Secretariat. CGIAR Research Program 2020 reviews: WHEAT. Rome: CAS Secretariat Evaluation Function. 2020. Available from: https://cas.cgiar.org/

12. Fischer RA, Byerlee D, Edmeades GO. Will yield increase continue to feed the world? Crop yields and global food security. Canberra: Australian Centre for International Agricultural Research. 2014.

13. IPCC. Pachauri RK, Reisinger A, editors. Contribution of Working Groups I, II and III to the Fourth Assessment Report of the Intergovernmental Panel on Climate Change., Climate Change 2007, Synthesis Report. Geneva: IPCC; 2007: 104.

14. Bita CE, Gerats T. Plant tolerance to high temperature in a changing environment: scientific fundamentals and production of heat stress-tolerant crops. Frontiers in Plant Sciences. 2013; 4: 273.

15. Buttar GS, Dhaliwal LK, Singh SP, Kingra PK. Climate change and agriculture mitigation and adaptations through agronomic practices. Asian Journal of environmental Science. 2012; 7(2): 239–244.

16. Suryavanshi P, Buttar G. Mitigating Terminal Heat Stress in Wheat. International Journal of Bio-resource and Stress Management. 2016. doi: 7.142-150.10.23910/IJBSM/2016.7.1.1333f.

17. Akther N, Islam R. Heat stress effects and management in wheat. A review. Agronomy Sustainable Development. 2017; 37: 37. doi: 10.1007/s13593-017-0443-9

18. Nahar K, Ahamed KU, Fujita M. Phenological variation and its relation with yield in several wheat (Triticum aestivum L.) cultivars under normal and late sowing mediated heat stress condition. Notulae Scientia Biologicae. 2010; 2: 51–56. doi: 10.15835/nsb234723

19. Pradhan GP, Prasad PVV. Evaluation of wheat chromosome translocation lines for high temperature stress tolerance at grain filling stage. PLoS One 10: e0116620. 2015. doi: 10.1371/journal.pone.0116620

20. Grant RF, Kimball BA, Conley MM, White JW, Wall GW, Ottman MJ. Controlled warming effects on wheat growth and yield: field measurements and modeling. Agron J. 2011; 103(6): 1742–1754. doi: 10.2134/agronj2011.0158

21. Joshi MA, Faridullah S, Kumar A. Effect of heat stress on crop phenology, yield and seed quality attributes of wheat (Triticum aestivum L.). J Agrometeorology. 2016; 18 (2): 206–215.

22. Bhatta MR, Sharma RC, Ortiz-Ferrara G. Wheat production and challenges in Nepal.: Reynolds MP, Pietragalla J, Braun HJ, editors. International Symposium on Wheat Yield Potential: Challenges to International Wheat Breeding, Mexico, D.F.: CIMMYT (International Center for Maize and Wheat Improvement). 2008.

23. Tripathi SC, Sayre KD, Kaul JN. Planting systems on lodging behavior yield components, and yield of irrigated spring bread. Wheat Crop Sci. 2005; 45: 1448–1455. doi: 10.2135/cropsci2003-714

24. Poudel MR, Ghimire S, Pandey MP, Dhakal KH, Thapa DB, Poudel HK. Evaluation of wheat genotypes under irrigated, heat stress and drought condition. Journal of Biology Today’s World. 2020; 9(1): 212.

25. Zampieri M, Ceglar A, Dentener F, Toreti A. Wheat yield loss attributable to heat waves, drought and water excess at the global, national and subnational scales. Environment Resesearch Letter. 2017; 12. doi: 10.1088/1748-9326/aa723b

26. Pradhan GP, Prasad PVV, Fritz AK, Kirkham MB, Gill BS. Effects of drought and high temperature stress on synthetic hexaploid wheat Funct. Plant Biol. 2012; 39: 190–8

27. Chapman SC, Chakraborty S, Dreccer MF, Howden SC. Plant adaptation to climate change-opportunities and priorities in breeding. Crop Pasture Sci. 2012; 63:251–268. doi: 10.1071/CP11303

28. Nezhadahmadi A, Prodhan ZH, Faruq G. Drought Tolerance in Wheat. Scientific World Journal. 2013. doi: 10.1155/2013/610721.

29. Jat ML, Jat RK, Singh P, Jat SL, Sidhu HS, Jat HS, Bijarniya D, Parihar CM, Gupta R. Predicting Yield and Stability Analysis of Wheat under Different Crop Management Systems across Agro-Ecosystems in India. American. Journal of Plant Sciences. 2017; 8: 1977–2012. Available from: http://www.scirp.org/journal/ajps

30. Vergas M, Crossa J, Eeuwijk FV, Sayre KD, Reynolds MP. Interpreting treatment ×environment interaction in agronomy trials. Agronomy Journals. 200; 93: 949–960.

31. Hamada AA, El-Hosary AA, El-Badawy MELM, Khalifa RM, El-Maghraby MA, Tageldin MHA, Arab SA. Estimation of phenotypic and genotypic stability for some wheat genotypes. Annals of Agric Sc Moshtohor. 2007; 45(1): 61–74

32. Zobel RW, Wright MJ, Gauch HG. Statistical analysis of a yield trial. Agron J. 1988; 80: 388–393.

33. Ding M, Tier B, Yan W, Wu HX, Powell MB, McRae TA. Application of GGE biplot analysis to evaluate Genotype (G), Environment (E), and G×E interaction on Pinus radiata: a case study. New Ze-aland Journal of Forestry Science. 2008; 38(1): 132–142. Available from: http://www.scionresearch.com/data/assets/pdf_file/0007/5596/NZJFS_38_12008_Ding_et_al_132-142.pdf

